# Calcium induced calcium release in proximity to hair cell BK channels revealed by PKA activation

**DOI:** 10.1101/2019.12.30.890558

**Authors:** Jun-ping Bai, Na Xue, Omolara Lawal, Anda Nyati, Joseph Santos-Sacchi, Dhasakumar Navaratnam

**Affiliations:** Department of Neurology, Yale School of Medicine, New Haven, CT, USA; Department of Surgery, Yale School of Medicine, New Haven, CT, USA; Department of Cell and Molecular Physiology, Yale School of Medicine, New Haven, CT, USA; Department of Neuroscience, Yale School of Medicine, New Haven, CT, USA; Department of Otolaryngology-Head and Neck Surgery, Shanghai Ninth People’s Hospital, Shanghai Jiaotong University School of Medicine, Shanghai, China; Undergraduate Program, Johns Hopkins University, Baltimore, MD, USA

## Abstract

Large conductance calcium-activated potassium (BK) channels play a critical role in electrical resonance, a mechanism of frequency selectivity in chicken hair cells. We determine that BK currents are dependent on inward flow of Ca^2+^, and intracellular buffering of Ca^2+^. Entry of Ca^2+^ is further amplified locally by Ca^2+^ induced Ca^2+^ release (CICR) in close proximity to plasma membrane BK channels. Ca^2+^ imaging reveals peripheral clusters of high concentrations of Ca^2+^ that are suprathreshold to that needed to activate BK channels. PKA activation increases BK currents likely by recruiting more BK channels due to spatial spread of high Ca^2+^ concentrations in turn from increasing CICR. STORM imaging confirms the presence of nanodomains with ryanodine and IP3 receptors in close proximity to the Slo subunit of BK channels. Together, these data require a rethinking of how electrical resonance is brought about and suggest effects of CICR in synaptic release. Both genders were included in this study.

## Introduction

Large conductance potassium channels (BK) play an essential role in hair cell physiology. In mammalian inner hair cells, these channels are the largest contributor to its outward current. In mammalian outer hair cells, these channels lie in proximity to nicotinic receptors and serve to set the resting membrane potential. In non-mammalian vertebrates, BK channels play an essential role in electrical resonance, a mechanism of frequency selectivity (Fuchs and Evans, 1990; Art et al., 1995; Fettiplace and Fuchs, 1999; Duncan and Fuchs, 2003).

In the best-studied example of turtles, electrical resonance is brought about by an interplay of an inward current through voltage-gated Ca^2+^ channels, and an outward current from large-conductance Ca^2+^ activated potassium (BK) channels (Fettiplace and Fuchs, 1999). These two channels lie in close proximity to one another and bring about oscillation in membrane potential (Roberts et al., 1990; Fettiplace and Fuchs, 1999). The frequency of membrane potential oscillation varies as a function of tonotopicity (Crawford and Fettiplace, 1981). In the turtle, this change in frequency is in turn brought about by variation in the number of channels and, more importantly, a change in the kinetics of the BK channel (Fettiplace and Fuchs, 1999). These data have been mainly corroborated in the chick auditory epithelium (Fettiplace and Fuchs, 1999; Duncan and Fuchs, 2003).

How might the changes in BK channel kinetics be brought about? The early promise of varying primary structure of the alpha subunit of the BK channel from changing alternative splicing along the tonotopic axis failed to explain the variation in channel kinetics (Jones et al., 1999; Ramanathan et al., 1999; Miranda-Rottmann et al., 2010). Changes in association with auxiliary proteins and changes in kinase activity along the tonotopic axis are two other mechanisms that could account for the alterations in the kinetic properties of the BK channel. Prior data have shown expression of KCNMB1 and KCNMB4 in the low-frequency end of the basilar papilla (Ramanathan et al., 2000; Bai et al., 2011), and, indeed, we demonstrated changes in CDK5 expression along the tonotopic axis (Bai et al., 2012b). Furthermore, higher PKA expression at the low-frequency end of the tonotopic axis is suggested by global gene expression analysis along the tonotopic axis (Frucht et al., 2011).

In this paper, we sought to determine how PKA activity affects BK channel kinetics in tall hair cells that receive principally afferent innervation. Our unexpected finding was that PKA recruited BK channels by inducing calcium-induced calcium release (CICR). Using super-resolution microscopy, we determine expression of clusters of both IP3 and ryanodine receptors along the plasma membrane of these cells in proximity to Slo, the alpha subunit of BK channels. These data have implications for the speed and amplification of feedback loops governing electrical tuning, and for synaptic vesicle release.

## Methods

All of the studies were done in accordance with the National Institutes of Health Guide for the Care and Use of Laboratory Animals, and the protocols were applied in compliance with the Yale University institutional review board guidelines.

### Electrophysiological recording

Whole-cell patch-clamp recordings were made of tall hair cells from freshly isolated basilar papilla from E21 chicks of either sex, when hearing is mature (Saunders et al., 1973; Fuchs and Sokolowski, 1990; Jones and Jones, 1995). Hair cells were exposed as previously described (Ricci et al., 2013). Recordings were obtained in the whole-cell configuration. Hair bundle heights were measured as a reference to its tonotopic location (Tilney et al., 1986). Cells were recorded in the presence of blockers of KCNQ (100 µM linopirdine) and SK channels (300 nM apamin), to isolate BK currents. The extracellular solution was as follows (in mM): 144 NaCl, 0.9 MgCl_2_, 1.3 CaCl_2_, 0.7 NaH_2_PO4, 10 HEPES, and 5 glucose, pH 7.4 and 300 mOsm. The composition of the pipette solution was (in mM): 120 K-gluconate, 20 KCl, 5 EGTA, 5 HEPES, 2.5 Na2ATP, 3.5 MgCl_2_, and 10 Na-phosphocreatine, pH 7.2 and 300 mOsm, and CaCl_2_ was added to reach the appropriate free Ca^2+^ concentration. The amount of total CaCl_2_ needed to obtain the desired free Ca^2+^ concentration was calculated with Max Chelator (https://somapp.ucdmc.ucdavis.edu/pharmacology/bers/maxchelator/webmaxc/webmaxcE.htm). Final free Ca^2+^ concentration was measured with a Ca^2+^ electrode (Thermo Electron, Beverly, MA) and confirmed our calculations. The extracellular solutions were delivered with the ALA QMM micromanifold perfusion system (ALA Scientific Instrument, Westbury, NY). Recordings were made at room temperature with an Axon 200B amplifier (Axon Instruments, Sunnyvale, CA). Command delivery and data collections were carried out with a Windows-based whole-cell voltage-clamp program, jClamp (Scisoft, Ridgefield, CT), using a Digidata 1322A interface (Axon Instruments). A standard protocol was adopted consisting of stepping the membrane potential from a holding potential of −80 mV to membrane potential 80 mV at 20-mV increments for 100 ms. The clock speed was set at 10 microseconds. Currents were digitized at 100 kHz and filtered at 5–10 kHz. Pipette resistance was ∼3-5 MΩ. Current-voltage (I-V) curves were obtained by measuring the averaged amplitude of currents at steady state after depolarization to various test voltages from holding the cells at −80 mV. Seal resistances for the recordings ranged from 0.5-2 GΩ, (mean 1 +/− 0.06 GΩ, median 0.8 GΩ). We corrected for junctional potentials owing to differences in Cl^-^ concentrations in the pipette and bath solutions. Correction for voltage errors due to the uncompensated series resistance was done offline. Similarly, leak currents were subtracted by estimating linear currents extrapolated from slopes at −95 to −80mV (corrected) where currents were linear.

V1/2 values were calculated from G-V curves. Conductance (G) was derived from Hodgkin and Huxley (Hodgkin and Huxley, 1952) and normalized (G/Gmax) to derive relative conductance to voltage relationships.

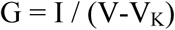

where G= conductance, V = step voltage, V_K_ = equilibrium potential for K^+^ (calculated to be −122mV), and I = current at that step voltage.

Each *G-V* curve was fitted with a Boltzmann function:

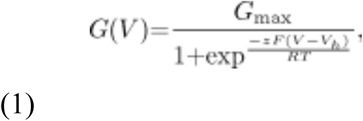

where Gmax is the fitted value for maximal conductance, Vh is the voltage of half maximal activation of conductance, and z reflects the net charge moved across the membrane during the transition from the closed to the open state. Data are reported as mean ± SE.

### Calcium imaging with confocal microscopy

Imaging was performed of chick hair cells from freshly isolated basilar papilla from E21 chicks as previously described for a chloride sensor developed in our lab (Zhong et al., 2014; Zhong et al., 2019). Chick hair cells were incubated with 1 µM Fluo-3-AM for 30 minutes at 22-24^°^C. Stock solutions of 1 mM Fluo-3-AM in DMSO were diluted to 1 µM in aquous solution. Typically, cells were incubated in the presence of artifical perilymph (in mM: 144 NaCl, 0.9 MgCl_2_, 1.3 CaCl_2_, 0.7 NaH_2_PO4, 10 HEPES, and 5 glucose, pH 7.4 and 300 mOsm). Perilymph containing 1 µM Fluo-3-AM was substituted with periplymph at the time of fluorescence measurement. In experiments with nominally 0 µM Ca^2+^, CaCl_2_ was removed and 2 mM EGTA added with the remaining constituents of the perilymph solution remaining constant. The papilla was mounted on a glass microtek dish under two insect pins, and Ca^2+^ signal was visualized while exciting at 488 nm using a Zeiss inverted spinning disc confocal microscope (Zeiss Observer Z1) with a 40X objective using 0.4 µm optical sections. For control experiments hair cells were incubated with medium containing specific concentrations of Ca^2+^ in the presence of the Ca^2+^ ionophore A23187 (1 µm) and 1 µM Fluo-3-AM for 30 minutes at 22-24^°^C (Dedkova et al., 2000). Where we measured effects of drugs in a time dependent manner, fluorescence from the same cells were tracked with the focal plane unchanged. Drift correction was applied to compensate for specimen drift. Image data were quantified with background correction using Zeiss Zen and Fiji software.

### Stochastic optical reconstruction microscopy (STORM) microscopy

Chick basilar papilla was labeled following protocol for super-resolution microscopy. In brief, freshly isolated basilar papillae were isolated and hair cells exposed by removal of the tectorial membrane following treatment with 0.5% collagenase for 4-5 minutes. Tissue was pre-extracted with 0.2% saponin followed by a fixation with 3% PFA and 0.1% glutaraldehyde. The tissue was reduced with 0.1% NaBH and labeled with primary (1:50) and secondary antibody (1:400, donkey antimouse Alexa 647 and donkey antirabbit Alexa 561) after blocking, with three washes of 3 minutes each between each step. The sample was post-fixed after antibody labeling with 4% PFA for 5 minutes. Freshly made imaging buffer containing glucose oxidase, catalase, mercaptoethanol, and MEA was added just before imaging. Super-resolution STORM images were obtained with the Bruker Vutara SR352 (Bruker Nano Surfaces, Salt Lake City, UT) with a 60x 1.2 NA objective and a 1 W 561 nm and 640 nm laser. Imaging beads confirmed that resolution was 20 nm in the *xy* plane and 50 nm in the *z*-direction. Calibration before experimentation was done by calculating the point spread function (PSF) in three dimensions using beads. Images were rendered and analyzed with Vutara’s SRX localization and visualization software (v6.2). Images were obtained in both planes simultaneously. The background was removed after the frames were obtained and particles identified on their brightness. Three-dimensional localization of the particles was based on 3D model function that was obtained from recorded bead datasets. The recorded fields are aligned automatically by computing the affine transformation between the pair of planes. Typically, we collected 5000 frames with each fluorophore using 20 µsec times. Data were analyzed using algorithms embedded in the Vutura software. These include the Crossed Nearest Neighbor algorithm and cluster identification. All chemicals were purchased from Sigma-Aldrich. Primary antibodies were as follows: mouse IgG2a monoclonal anti-BK channel α subunit antibody (BD Labs) (Surguchev et al., 2012), anti-BK channel α subunit polyclonal antibody (APC021) (Alomone labs, Jerusalem, Israel) (Purcell et al., 2011); IgG1 monoclonal anti-ryanodine receptor antibody (clone 34C, Developmental Studies Hybridoma Bank, University of Iowa, Iowa City, Iowa; this antibody detects all RyR isoforms in mouse tissue (Irie and Trussell, 2017); rabbit polyclonal anti IP3R2 antibody (Alomone labs, Jerusalem, Israel(Tadevosyan et al., 2017; Sabourin et al., 2018). Secondary antibodies were as follows: AF 568 goat anti-mouse IgG, AF 568 goat anti-rabbit IgG, Alexa Fluor 647 goat anti-rabbit IgG, and Alexa Fluor 647 goat anti-mouse IgG (Jackson labs, Maine). We used mouse monoclonal Slo antibody with the polyclonal rabbit IP3 antibody (and corresponding conjugated secondary antibodies) to detect co-localization of these two proteins. For experiments to detect ryanodine receptor and Slo protein we used the Slo polyclonal rabbit antibody and ryanodine monoclonal mouse antibody together with the corresponding secondary antibodies. All primary antibodies were used at a concentration of 1µg/ml. All secondary antibodies were used at a 1:200 dilution. These antibodies have all been previously validated.

## Results

### Hair cells possess a BK current that is sensitive to entry of extracellular Ca^2+^, and intracellular Ca^2+^ buffering

Current recordings of chick hair cells from the neural edge, 20-25% of the distance of the basilar papilla from the apical end, were obtained under whole-cell voltage-clamp conditions. We confirmed the location of these hair cells using stereociliary height (Tilney et al., 1986). As previously demonstrated (Fuchs et al., 1988; Fuchs and Evans, 1990) (Fuchs and Sokolowski, 1990), these cells demonstrated a large outward current with 140mM KCl in the pipette and 140mM NaCl in the bath (Figure 1A). Consistent with previous experimental data, the majority of the current was carried by a large-conductance Ca2+ activated K channel (Fuchs et al., 1988; Fuchs and Evans, 1990; Fuchs and Sokolowski, 1990; Duncan and Fuchs, 2003). The outward current showed rapid rates of activation that is a hallmark of BK currents. Consistent with it being a BK current, it was blocked by extracellular TEA (20mM), partially blocked by 100 µM Penitrem A, an incomplete blocker of BK channels, and insensitive to 5mM 4-AP, a blocker of voltage-gated potassium channels (**Figure 1A-I**). The bath also contained 100 µM linopirdine, 300 nM apamin and 50 µM PPADS, blockers of KCNQ and SK channels and P2 purinergic receptors respectively, the other sources of outward currents in these cells. As previously reported, these BK current are insensitive to charybdotoxin and iberiotoxin owing to the high expression of the beta4 (KCNMB4) subunit that confers resistance to these blockers (Reinhart et al., 1989; Brenner et al., 2000; Meera et al., 2000; Brenner et al., 2005; Gan et al., 2008; Bai et al., 2012a).

**Figure 1.**
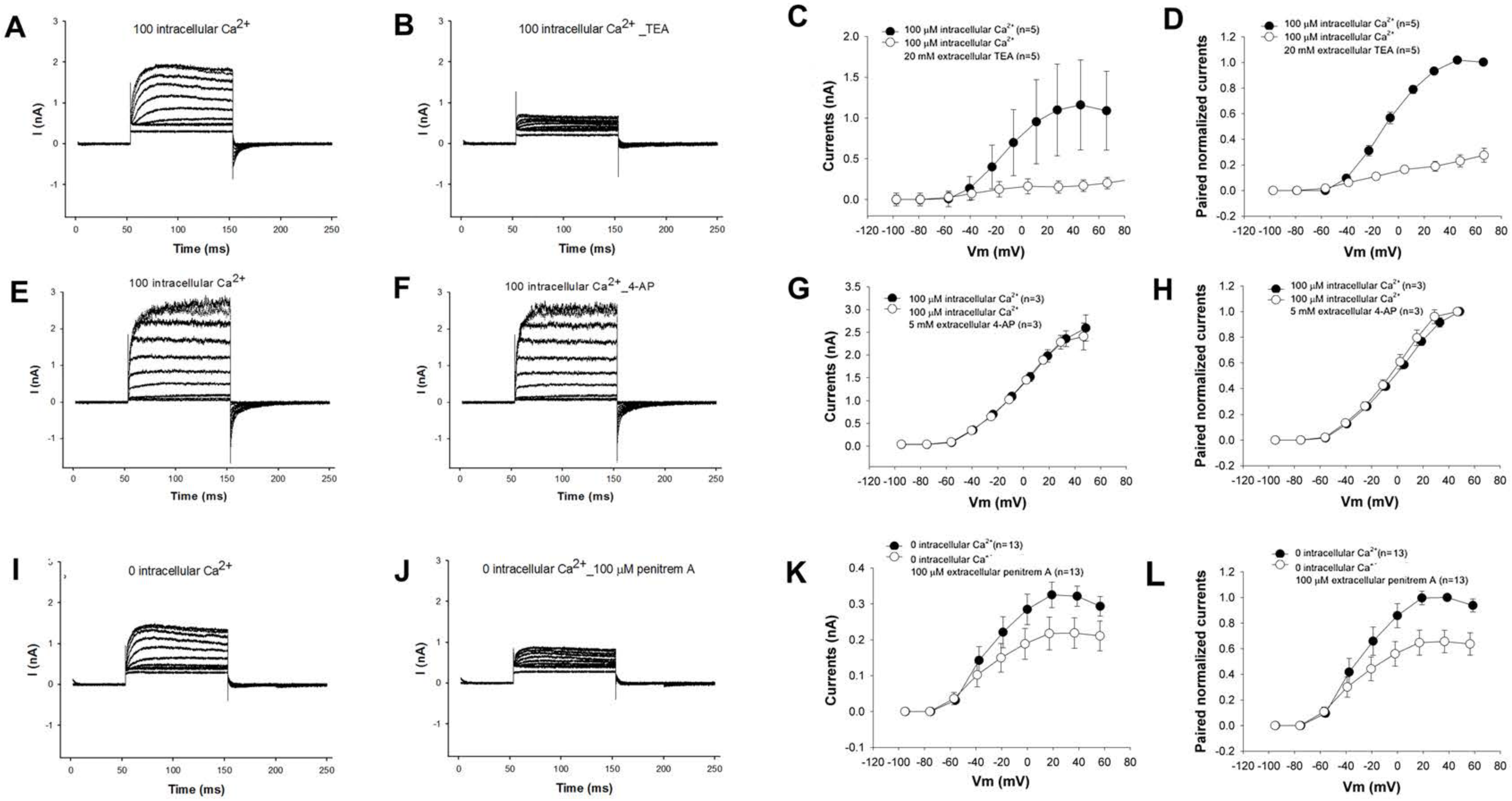
Hair cells have a large outward potassium current. **A-D.** The outward current is reduced by the application of extracellular 20mm TEA. Shown are individual representative current recordings of a cell under voltage clamp before (**A**) and after (**B**) perfusion with extracellular TEA. Averaged I-V relationships (**C**) and paired averaged currents normalized to preperfusion currents (**D**) of cells perfused with TEA shows a marked reduction in the size of the current with TEA perfusion. (**C**). **E-H.** The size of the current is unaffected by 100 µM 4AP. Shown are representative current recordings of an individual cell before (**E**) and after perfusion (**F**) with 4-AP. Averaged I-V relationships (**G**) and paired averaged currents normalized to preperfusion currents (**H**) of cells perfused with 4AP show no change in the size of the current with 4AP perfusion.**I-L**. The size of the current is partially reduced by 100 µM of Penitrem A, a partial inhibitor of BK channels. Representative current recordings of cells under voltage clamp before (**I**) and after (**J**) perfusion with 100 µM Penitrum A. Averaged I-V relationships (**K**) and paired averaged currents normalized to preperfusion currents (**L**) of cells perfused with Penitrum A show a moderate reduction in the size of the current with Penitrum A perfusion.

The size of the outward current and its voltage sensitivity was dependent on the inward flow of Ca^2+^. The size of the current decreased and showed a rightward shift in its voltage-current relationship when voltage-gated Ca^2+^ channels were blocked with 100 µM CdCl_2_ (**Figure 2A-E**). V_½_ (calculated from G-V curves) was shifted significantly from −37 mV to −2 mV after perfusion with CdCl_2_. There was a similar significant shift in V_1/2_ from −21 mV to 32mV (again calculated from G-V curves) and a reduction in the size of the current (**Figure 2 F-J**) when extracellular Ca^2+^ was chelated (with 3 mM EGTA).

**Figure 2.**
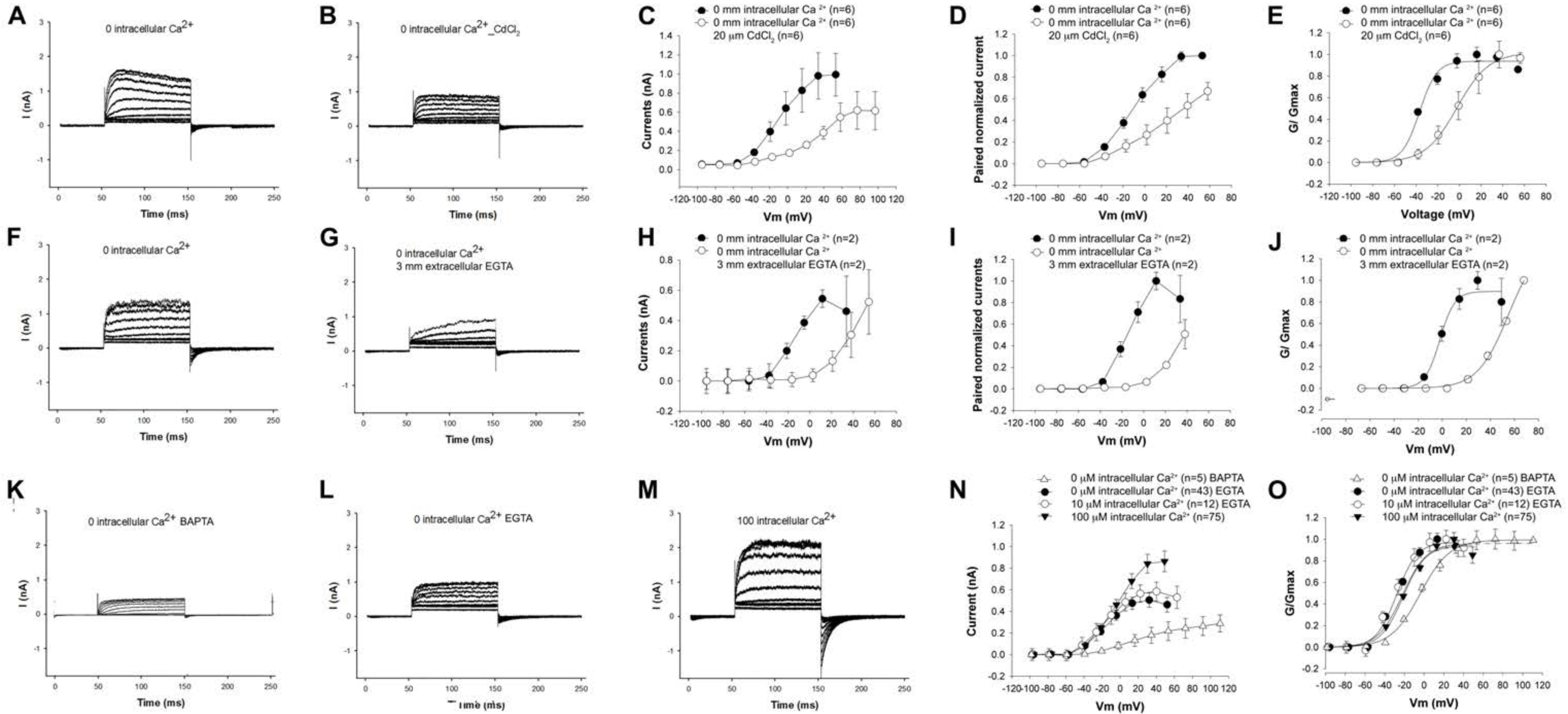
Hair cell BK currents are dependent on extracellular Ca^2+^ and intracellular Ca^2+^ buffer that have similar effects. **A-E** Extracellular CdCl_2_ reduces the size of the outward current. Representative current recordings of hair cells under voltage clamp before (**A**) and after (**B**) perfusion of 20 µM CdCl_2_ into the extracellular bath solution. Averaged I-V relationships (**C**) and paired averaged currents normalized to preperfusion currents (**D**) of cells perfused with CdCl_2_ shows a reduction in the size of the current at comparable voltages with CdCl_2_ perfusion. Currents are also significantly shifted in a depolarizing direction after perfusion (**E**). **F-J**. A similar effect is seen with reducing extracellular Ca^2+^. Shown are representative current recordings of hair cells under voltage clamp before (**F**) and after (**G**) perfusing with bath solution containing 3 mM EGTA. Averaged currents (**H**) and averaged paired currents normalized to currents before perfusion (**I**) show a reduction in the size of the currents with EGTA perfusion. Currents show a significant depolarizing shift in its current-voltage relationship after reducing the extracellular Ca^2+^ concentration **(J). K-O.** Specific buffers and intracellular Ca^2+^ variably affect the size of the current and its I-V relationship. Shown are representative current recordings of hair cells under voltage clamp with nominally 0 µM Ca^2+^ buffered with BAPTA in the pipette (**K**), nominally 0 µM Ca^2+^ buffered with EGTA in the pipette (**L**) and 100 µM Ca^2+^ in the pipette (**M**). Averaged currents (**N**) show a reduction in the size of the currents with reducing intracellular Ca^2+^ that was especially marked with greater spatial buffering afforded by BAPTA. Currents show a depolarizing shift in its current-voltage relationship with greater spatial buffering provided with BAPTA (**O**). In contrast, increasing pipette Ca^2+^ concentrations with EGTA as the buffer increased the size of the current without affecting the I-V relationship.

The size and voltage dependence of the current is dependent on the intracellular Ca^2+^ buffer (**Figure 2 K-O**). The voltage dependence was significantly shifted in a depolarizing direction when using BAPTA as the intracellular buffer (with nominally 0 µM Ca^2+^) compared to using EGTA as the intracellular buffer (again, with nominally 0 µM Ca^2+^). V_½_ was shifted significantly from −27 mV to −4 mV with EGTA and BAPTA as the intracellular buffers, respectively (V_½_ was estimated from G-V curves). The size of the outward current was also significantly reduced when BAPTA (nominally 0 µM Ca^2+^) was used as the intracellular buffer when compared to using intracellular EGTA (nominally 0 µM Ca^2+^). These data argue that spatial buffering by BAPTA significantly attenuates influx of Ca^2+^ in proximity to BK channels. It has been estimated that buffering by BAPTA limits the spread of Ca^2+^ to 20-50 nm (nanodomains). In contrast, owing to the slower on rate of EGTA spatial buffering is limited to micro-domains (> 50nm) (Heidelberger et al., 1994; Neher, 1998; Augustine et al., 2003).

Increasing intracellular Ca^2+^ in the presence of EGTA from 0 to 10 µM resulted in an increase in the size of the current (**Figure 2 K-O**). The size of the current was further increased in the presence of 100 µM Ca^2+^. However, these increases in the size of the current were accompanied by a minimal shift in voltage sensitivity with V_1/2_ ranging from −29mV to −21mV (**Figure 2 K-O**). Along with the data from recordings in the presence of BAPTA, these data argue that the local concentration of Ca^2+^ in proximity to BK channels from inward flow of Ca^2+^ is spatially buffered by BAPTA. They also argue that the local concentration of Ca^2+^ around BK channels is saturating in the presence of EGTA with nominally 0 µM and higher Ca^2+^ (when spatial buffering is more limited). Finally, these data suggest that spatial buffering of intracellular Ca^2+^ is a possible mechanism of recruiting BK channels.

### Perfusion with activators of PKA increased the size of the outward current

Perfusing cells with 100 µM forskolin increased the size of the outward current by twofold (**Figure 3A-C**), an effect similar to increasing intracellular (pipette) Ca^2+^ (Figure 2). The effect of forskolin was dependent on the concentration of intracellular Ca^2+^. Thus, the forskolin-induced increase in outward current was evident with 100 µM intracellular (pipette) Ca^2+^ with EGTA (**Figure 3A-E**). In contrast, there was a no increase in the size of the current in the presence of nominally 0 µM intracellular (pipette) Ca^2+^ with EGTA (**Figure 3F-J**). The effects of forskolin in the presence of 10 µM Ca^2+^ was intermediate (data not shown). V_1/2_ was affected, shifting a significant −5 mV to –17 mV with forskolin (with 100 µM pipette Ca^2+^). Confirming a basal activation of PKA in hair cells contributing to BK channel activity, with 100 µM Ca^2+^ in the pipette, perfusion of the PKA inhibitor H-89 reduced the size of the outward current by approximately 1/3 (**Figure 3K-O**). This reduction in the size of the current was not accompanied by a change in the voltage-current relationship (**Figure 3K-O**). V_½_ was shifted by a non-significant −18 mV to −15 mV after perfusion with H-89. Together, these data suggest that PKA increases BK channel activity that is dependent on intracellular Ca^2+^ concentration and that hair cells have a basal level of PKA activity.

**Figure 3.**
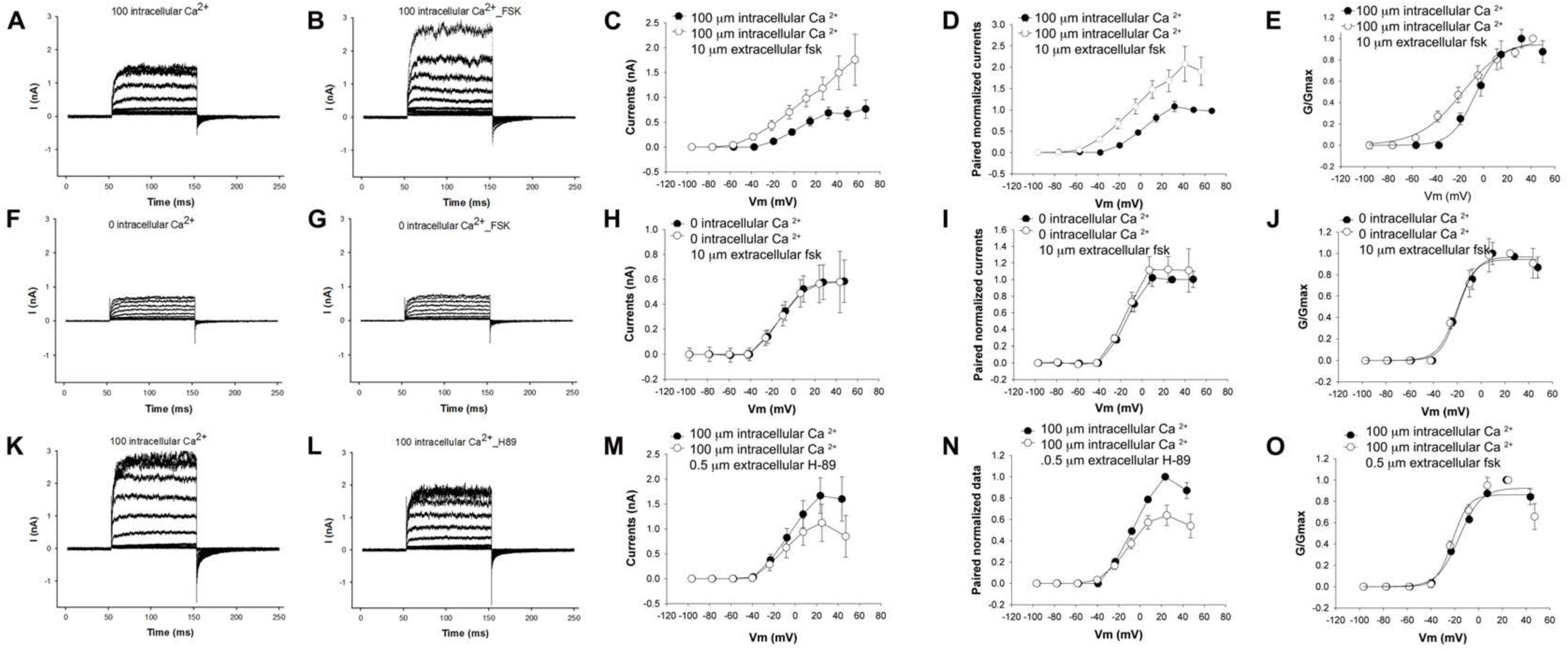
Forskolin increased the size of the outward current in a Ca^2+^ dependent manner. **A-C.** Representative current recordings of hair cells before (**A**) and after (**B**) perfusion with 100 µM forskolin with the pipette containing 100 µM Ca^2+^. The size of the outward current increased twofold when hair cells were perfused with 100 µM forskolin evidenced in averaged currents (**C**), and averages of pre and post perfusion paired currents normalized to pre perfusion currents (**D**). Currents also show a significant hyperpolarizing shift in voltage sensitivity (**E**). **F-K.** In contrast, perfusion of hair cells with 100 µM forskolin with nominally 0 µM pipette Ca^2+^ (with EGTA as the intracellular buffer) did not cause an increase in the size of the current or a change in its current-voltage relationship. Shown are representative current recordings of hair cells under voltage clamp before (**F**) and after (**G**) perfusion with 100 µM forskolin. The averaged I-V relationship of cells before and after perfusion with forskolin compared to pre perfusion (**H**) and paired averaged currents from individual cells normalized to values before perfusion (**I**) confirm an absence of an effect on the size of the current. G-V curves (**J**) to elicit voltage relationships do not show an effect on its I-V relationship. **K-O.** Perfusion of hair cells with 0.5 µM H-89, a well-known PKA blocker resulted in a reduction in the size of the current. Representative current recordings of cells under voltage clamp before (**K**) and after (**L**) perfusion with H-89 are shown. The averaged I-V relationships of cells before and after perfusion with H-89 (**M**) and averaged currents from individual cells pre and post perfusion normalized to values before perfusion (**N**) confirm a decrease in the size of the currents with H-89. G-V curves (**O**) to elicit voltage relationships show a non-significant change in voltage sensitivity.

### The effects of forskolin on the outward current are due to Ca^2+^-induced Ca^2+^ release

How does PKA increase BK channel activity while minimally affecting V_1/2_? Direct effects on the channel bearing the STREX exon (the dominant exon in these hair cells) would be predicted to shift V_1/2_ in a depolarizing direction (Tian et al., 2001; Frucht et al., 2011). Since forskolin effects on hair cell outward current were dependent on pipette Ca^2+^ concentration, Ca^2+^-induced Ca^2+^ release was suggested as a likely mechanism. To test this possibility, we treated cells with an inhibitor of ryanodine receptors. Currents from hair cells under voltage clamp were recorded in the presence of 100 µM pipette Ca^2+^ and treated with 10 µM dantrolene, a potent blocker of ryanodine receptors (**Figure 4F-J**). There was a significant reduction in the size of the current.

**Figure 4.**
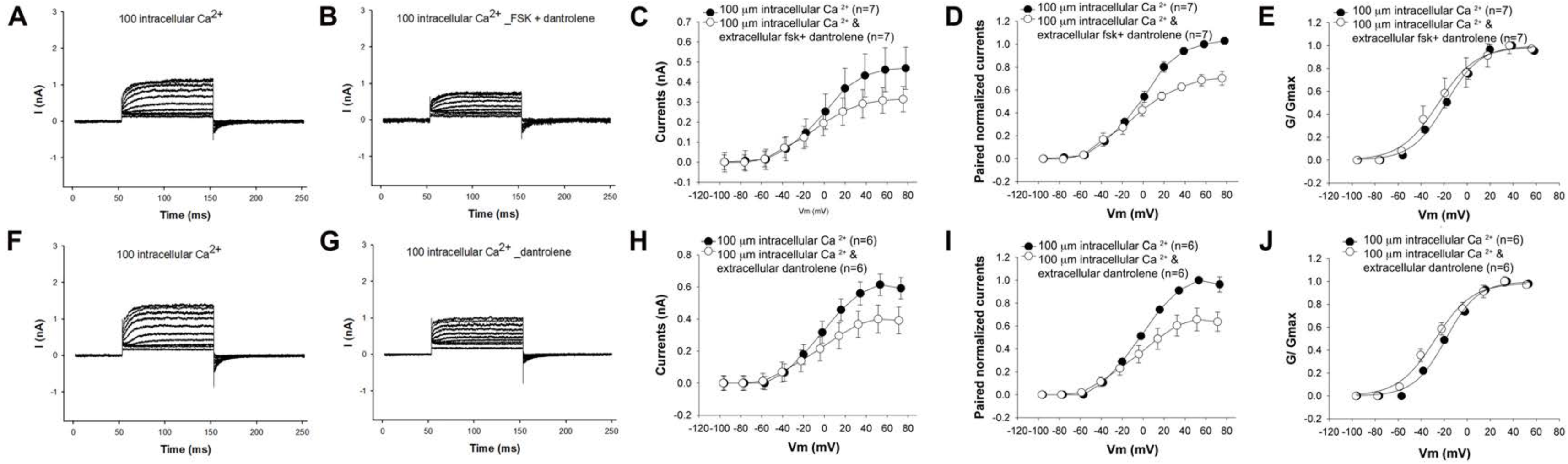
Blocking ryanodine receptors prevents a forskolin-induced increase in the outward current **A-E.** The IV relationship of the outward current before and after perfusion with 100 µM forskolin and 1 µM Dantrolene shows a decrease in the size of the current with no change in the IV relationship. The pipette contained 100 µM Ca^2+^. Shown are representative current recordings of cells under voltage clamp before (**A**) and after (**B**) perfusion with forskolin and dantrolene. Averaged currents (**C**) and averaged currents of pairs of currents before and after perfusion normalized to preperfusion currents (**D**) show an approximately 1/3 reduction in current in cells treated with both dantrolene and forskolin. G-V curves (**E**) show no change in voltage sensitivity. **F-J.** A similar reduction in the size of the current with no change in the IV relationship is seen when hair cells are perfused with 1 µM Dantrolene alone without forskolin. Here too, the pipette contained 100 µM Ca^2+^. Shown are representative current recordings of cells under voltage clamp before (**F**) and after (**G**) perfusion with dantrolene. The averaged currents (**H**) and averaged currents of pairs of currents before and after perfusion normalized to pre-pefusion currents (**I)** show a reduction in the size of the current after perfusion with dantrolene. Here too, the reduction in the size of the current was approximately 1/3 indicating that the addition of forskolin had no additional effect in the presence of dantrolene. G-V curves (**J**) show no change in voltage sensitivity.

The reduction in the size of the current was also not accompanied by a change in its voltage dependence. V_1/2_ shifted in a nonsignificant fashion from −20 mV to −29 mV. Separately, cells treated with 10 µM dantrolene and 100 µM forskolin showed a similar reduction in the size of the outward current (**Figure 4A-E**). Here too V_1/2_ shifted from −16 mV to a statistically insignificant −26 mV after treatment.

We also tested the ability of inhibitors of IP3 receptors (ITPRs) to block the effects of forskolin. Similar to dantrolene, cells treated with the IP3 receptor antagonist 100 µM 2-APB (2-aminoethoxydiphenyl borate), show a similar reduction in the size of the outward current (**Figure 5F-H**). The voltage sensitivity shifted in a depolarizing direction with V_1/2_ shifting from −21 mV to −12 mV. As with dantrolene and forskolin, hair cells separately treated with 100 µM 2-APB and 100 µM forskolin showed a decrease in the size of the outward current (**Figure 5A-E**). Unlike dantrolene however, there was a greater reduction in the size of the current when 2-APB was combined with forskolin (when compared to 2-APB alone). Currents were 40% of preperfusion values with 2APB and forskolin, in contrast to the effects of 2APB alone where currents were 60% of preperfusion values. Moreover, there was a significant hyperpolarizing shift in the V_1/2_ from −36 mV to −6 mV. We believe these effects on the size of the current and its I-V relationship to represent two causes; a reduction in local Ca^2+^ concentrations by preventing CICR, and a directly inhibitory effect of PKA on the Slo channel containing the STREX exon in the absence of local release of Ca^2+^ (Ramanathan et al., 2000; Chen et al., 2005; Frucht et al., 2011). Although these experiments were not specifically designed to address the concern of effects at a hair cells operating voltage, we noted that IP3 receptor blocking had a significant effect on the size of the outward current at a hair cells operating voltage, estimated to be −50 mV, in contrast to block of ryanodine receptors.

**Figure 5.**
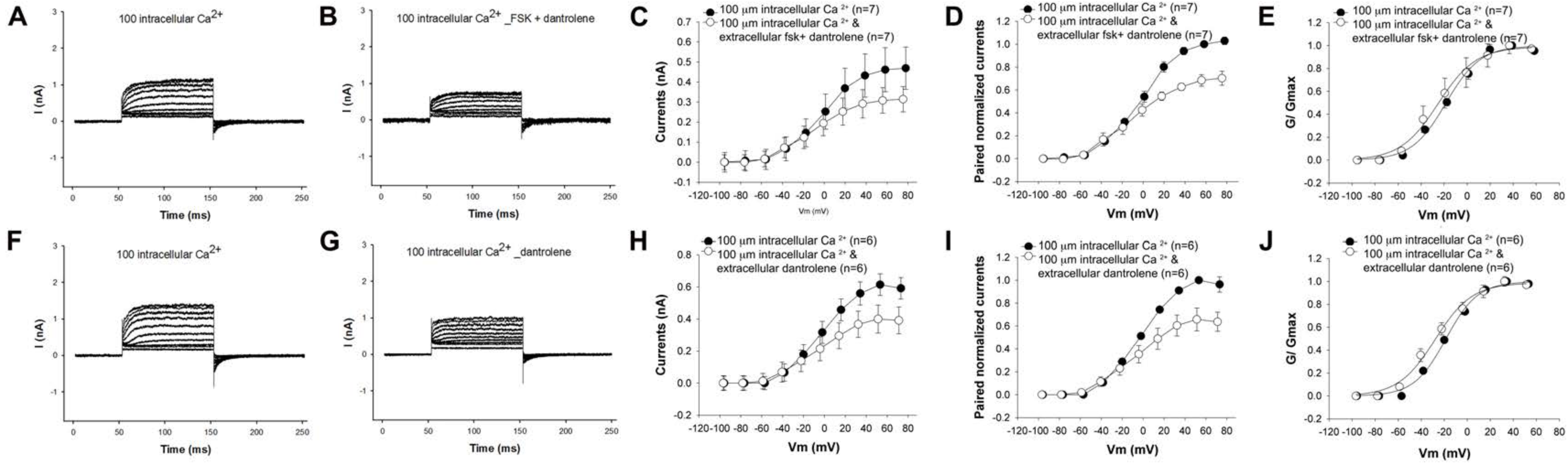
Blocking IP3 receptors prevents a forskolin-induced increase in the outward current. **A-E.** The IV relationship of the outward current before and after perfusion with 100 µM forskolin and 100 µM 2-APB shows a larger decrease in the size of the current. The pipette contained 100 µM Ca^2+^. Shown are representative current recordings of cells under voltage clamp before (**A**) and after (**B**) perfusion with forskolin and dantrolene. Averaged currents (**C**) and averages of paired currents before and after perfusion normalized to preperfusion currents (averaged currents of pairs of currents before and after perfusion **(D))** demonstrate current values to be 40% of the current before perfusion with 2-APB and forskolin. G-V curves (**E**) show a significant shift in voltage sensitivity in cells treated with forskolin and 2-APB. **F-J.** A smaller reduction in the size of the current with a change in the IV relationship is seen when hair cells are perfused with 100 µM 2-APB alone without forskolin. Here too, the pipette contained 100 µM Ca^2+^. Shown are representative current recordings of cells under voltage clamp before (**F**) and after (**G**) perfusion with 100 µM 2-APB. Averaged currents (**H**) and averages of paired currents before and after perfusion normalized to preperfusion currents (averaged currents of pairs of currents before and after perfusion (**I**) show a smaller 1/3 decrease in the size of the current when cells were perfused with 2-APB alone. G-V curves (**J**) show a small shift in voltage sensitivity after perfusion with 2-APB.

Together, these data confirm that PKA increases hair cell Ca^2+^ concentration in proximity to BK channels by a calcium-induced calcium release (CICR) mechanism, with inhibition of IP3 receptors having a bigger effect.

### Ca^2+^ imaging reveals clusters of Ca^2+^signal in the periphery of hair cells that is dependent on CICR

To confirm Ca^2+^ influx and its effects, we imaged hair cells loaded with the Ca^2+^sensor dye Fluor-3-AM. We noted a significant increase in the Fluor-3 signal when the cells were incubated with perilymph that contains 1.3 mM Ca^2+^. The signal was most notable along the periphery of the cell in axial sections when the cell was viewed end-on from above (**Figure 6A**). In cells viewed laterally, there was a significant increase in signal at the periphery of that was weighted to the lower half of the cell (**Figure 6C**). In contrast, cells kept in nominally 0 µM extracellular Ca^2+^ showed no peripheral increase in Ca^2+^signal (**Figure 6B**).

**Figure 6.**
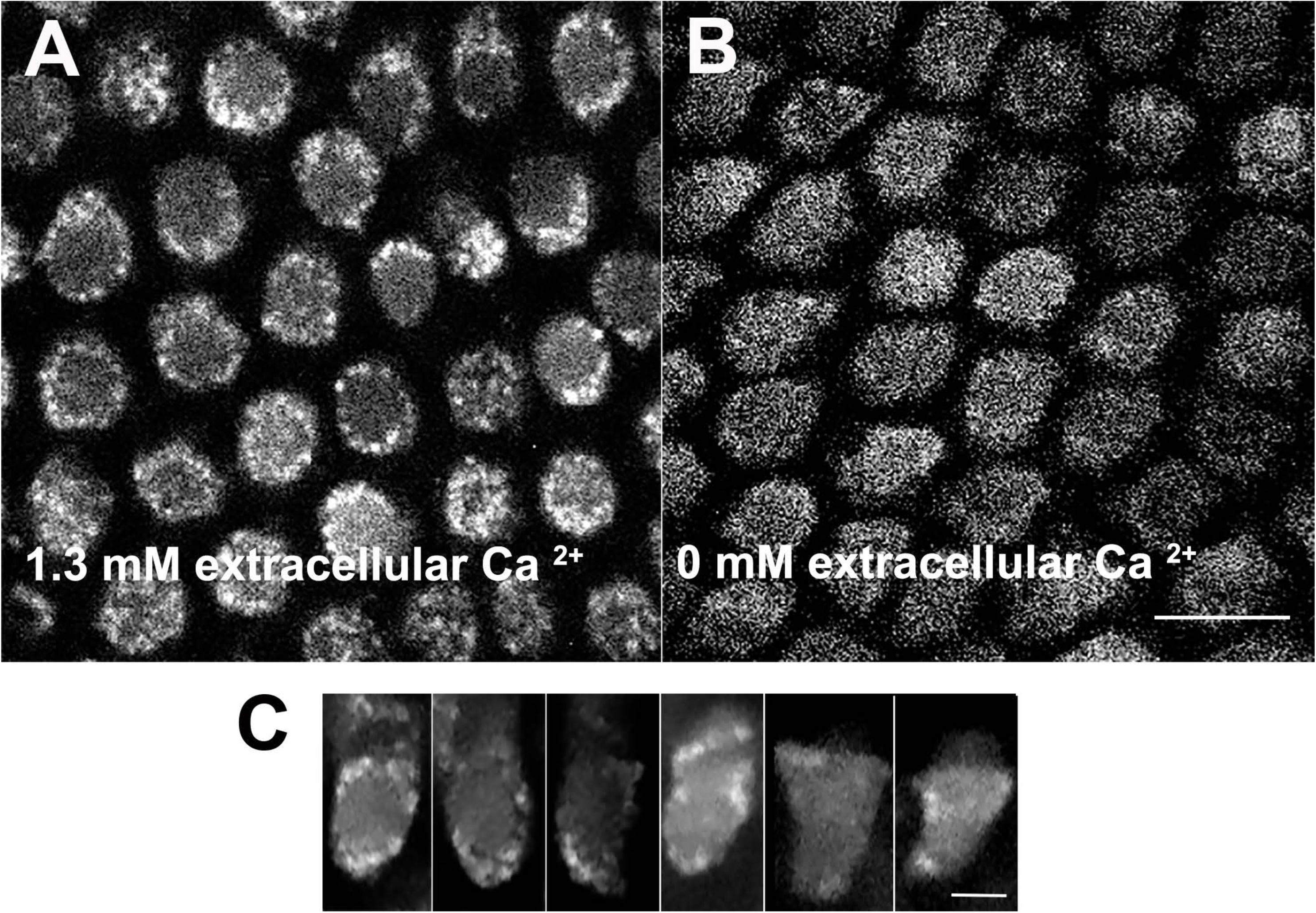
In artificial perilymph, hair cells show high concentrations of peripheral Ca^2+^ in clusters. **A.** Hair cells viewed end-on from above using confocal microscopy show high concentrations of peripheral Ca^2+^. Ca^2+^ was detected after incubating cells in 1 µM Fluor-3-AM in perilymph containing 1.3 mM Ca^2+^. These high concentrations of Ca^2+^ are not uniformly spread along the periphery and are clustered. **B.** Hair cells incubated with 1 µM Fluor-3-AM in perilymph containing nominally 0 µM Ca^2+^ for 30 minutes at room temperature are viewed end-on from above and show absent peripheral concentration of Ca^2+^ signal. Scale bar = 10 µm. **C.** Hair cells in basilar papillae incubated with 1 µM Fluor-3-AM in perilymph for 30 minutes at room temperature viewed side on show high concentrations of peripheral Ca^2+^. Here too, the Ca^2+^ signal is clustered. Also, note increased signal in stereocilia and in the region of the cuticular plate. Scale bar = 5 µm.

In contrast to the peripheral accumulation of signal in hair cells incubated with perilymph alone, the addition of inhibitors of both IP3 receptors (10 µM 2-APB) and ryanodine (100 µM dantrolene) resulted in a marked reduction in the intensity of Ca^2+^ signal (**Figure 7**). These findings were reflected in the gradient in peripheral Ca^2+^ signal that was significantly attenuated in the presence of these two inhibitors (**Figure 7**). We conclude that peripheral Ca^2+^ signal in hair cells is increased by physiological concentrations of extracellular Ca^2+^ that in turn induces local CICR.

**Figure 7.**
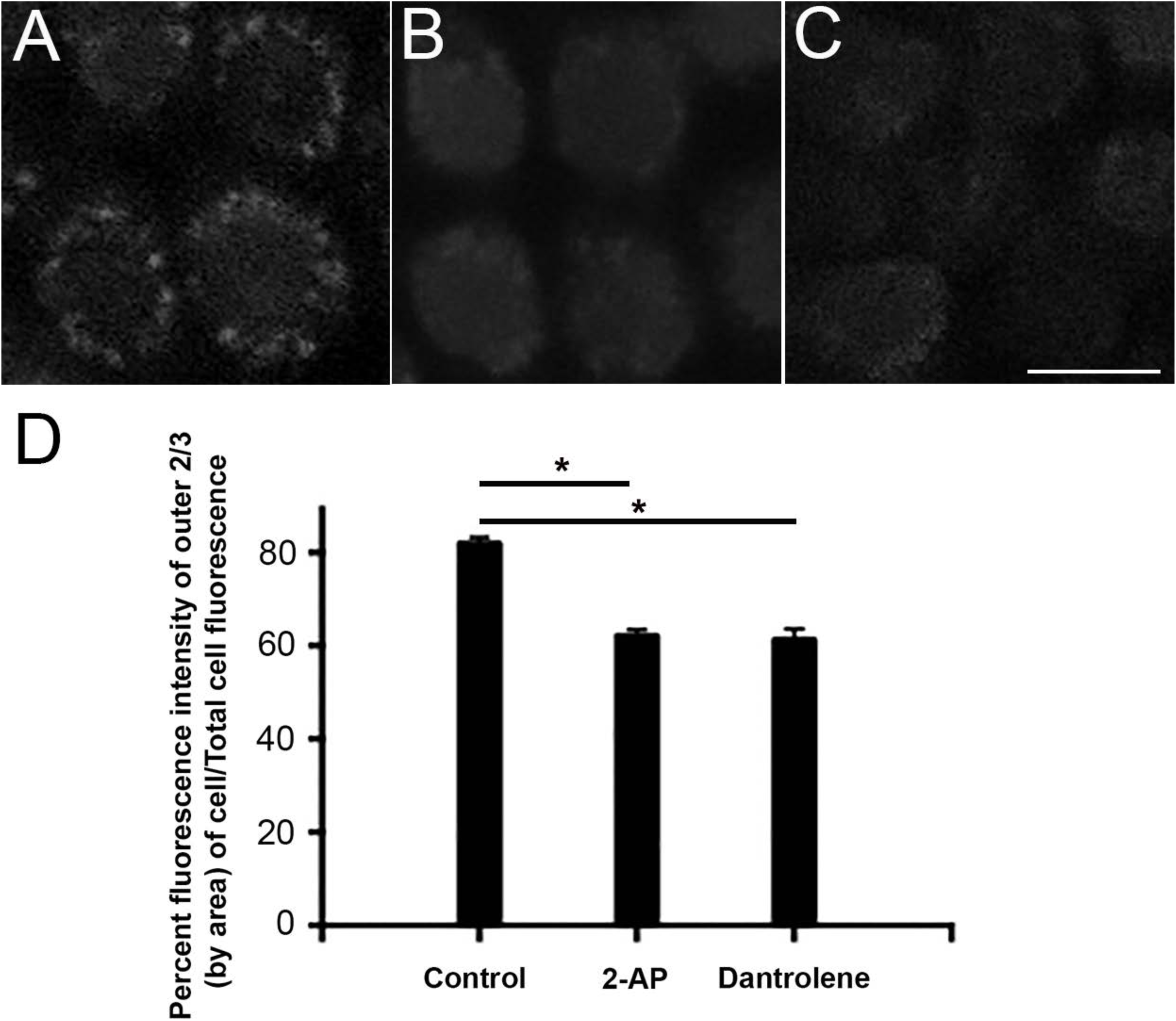
In perilymph, hair cells do not show high concentrations of peripheral Ca^2+^ in the presence of inhibitors of CICR. **A.** Control cells incubated in 1 µM Fluor-3-AM in perilymph (1.3 mM Ca^2+^) for 30 minutes shows a peripheral accumulation of Ca^2+^ along the periphery of the cell. Data were acquired using a spinning disc confocal microscope with the cells viewed end on from above. **B, C.** Addition of 100 µM 2-APB (B) or 1 µM dantrolene (C) both prevent an increase in the peripheral accumulation of Ca^2+^**. D.** These effects of inhibitors of CICR were quantified. Shown is the cumulative Ca^2+^ signal in the peripheral 2/3 of the cell viewed end-on. In the presence of inhibitors of CICR, there is a significant reduction in the accumulation of peripheral Ca^2+^ signal as a percentage of the signal in the entire cell (P<0.01, one-way ANOVA). Scale bar = 5 µm.

### 8-br-cAMP increases the local Ca^2+^ concentration, particularly at the periphery of the cell

We determined the effects of raising cAMP levels while monitoring intracellular Ca^2+^. **Figure 8** shows the effects of 100 µM 8-br-cAMP, the cell-permeable analog of cAMP that activates PKA, on intracellular Ca^2+^ concentration. We note a spike in Ca^2+^ concentration that followed treatment with 8-br-cAMP (**Figure 8**). The increase in Ca^2+^ signal was most notable along the periphery of the cell. In contrast, cells pretreated with 100 µM dantrolene and 100 µM 2-APB showed minimal to no increase in the size of the Ca^2+^ signal. Since the Ca^2+^ signal was significantly attenuated, and particularly along the periphery of the cell, by treating with IP3 and ryanodine antagonists, we used the Ca^2+^ signal in the entire cell for these comparisons. These data confirm that PKA activation increases peripheral Ca^2+^ concentrations by CICR.

**Figure 8.**
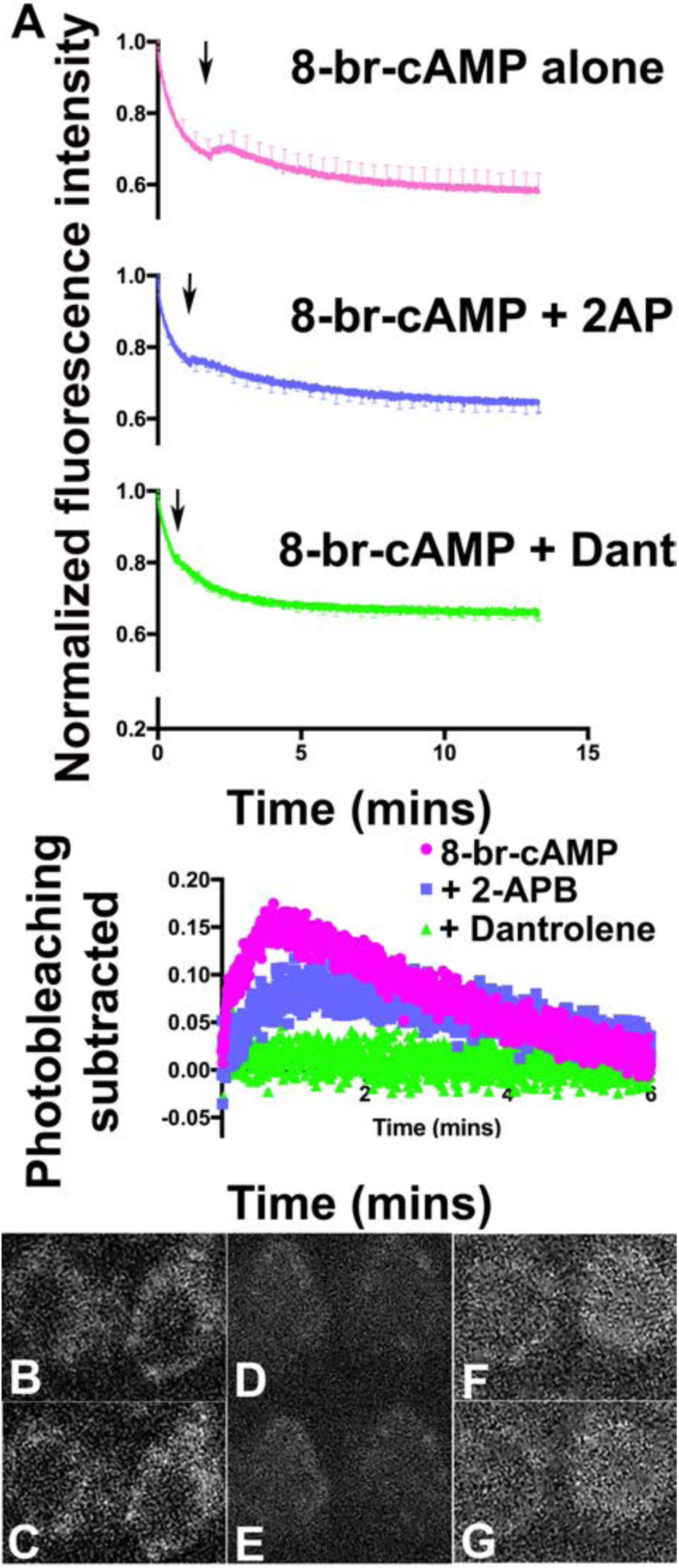
Addition of 8-br-cAMP increases Ca^2+^ signal in hair cells that is attenuated in the presence of inhibitors of CICR **A.** Shown are Ca^2+^ fluorescence signal (detected after incubating cells in 1 µM Fluor-3-AM in perilymph containing 1.3 mM Ca^2+^ for 30 minutes) in hair cells as a function of time. The addition of 100 µM 8-br-cAMP (arrow) resulted in an increase in Ca^2+^ in the cell above the exponential decay in fluorescence due to photobleaching. In contrast, the presence of 2-APB and dantrolene (both inhibitors of CICR) attenuated (2-APB) and eliminated (dantrolene) the increase in Ca^2+^ signal induced by 100 µM 8-br-cAMP. In order to make comparable measurements, Ca^2+^ signal in the entire cell was measured, since inhibitors of CICR caused a significant reduction in peripheral Ca^2+^ signal. The lowest panel shows the increase in Ca^2+^ signal from which the exponential decay from photobleaching was subtracted. In this panel, the fluorescence signal was synchronized to the point at which 8-br-cAMP was added. Error bars are SEM. **B,C,D,E,F,G.** Ca^2+^ imaging of individual hair cells before (B - 8-br-cAMP alone, D – 8-br-cAMP plus 2-APB, F – 8-br-cAMP plus Dantrolene) and after (C - 8-br-cAMP alone, E – 8-br-cAMP plus 2-APB, G - 8-br-cAMP plus Dantrolene) treatment with 8-Br cAMP. Cells were preloaded with 1 µM Fluor3 and the respective inhibitors for 30 minutes at RT C in perilymph containing 1.3 mM Ca^2+^.

### We made estimates of Ca^2+^ concentration in hair cells incubated in bath solution that approximated that of perilymph

Our cumulative data suggest a high concentration of Ca^2+^ in proximity to BK channels in the experimental conditions we used for our electrophysiological recordings. Prior work has demonstrated BK channels to lie in proximity to VGCCs at the periphery of the cell (Roberts et al., 1990; Issa and Hudspeth, 1994; Samaranayake et al., 2004). We sought to determine the concentration of Ca^2+^ along the periphery of the cell in the presence of perilymph. For these experiments, we calibrated the Ca^2+^ fluorescence by using the Ca^2+^ ionophore A23187 and incubated cells in different concentrations of external Ca^2+^ for 30 minutes before measuring Ca^2+^ signal (Dedkova et al., 2000). In **Figure 9**, we determine the concentration of Ca^2+^ along the periphery of hair cells incubated in perilymph to be in excess of 100 µM. These data are in broad agreement with our electrophysiological data suggesting local concentrations of Ca^2+^ in excess of that required to activate BK channels.

**Figure 9.**
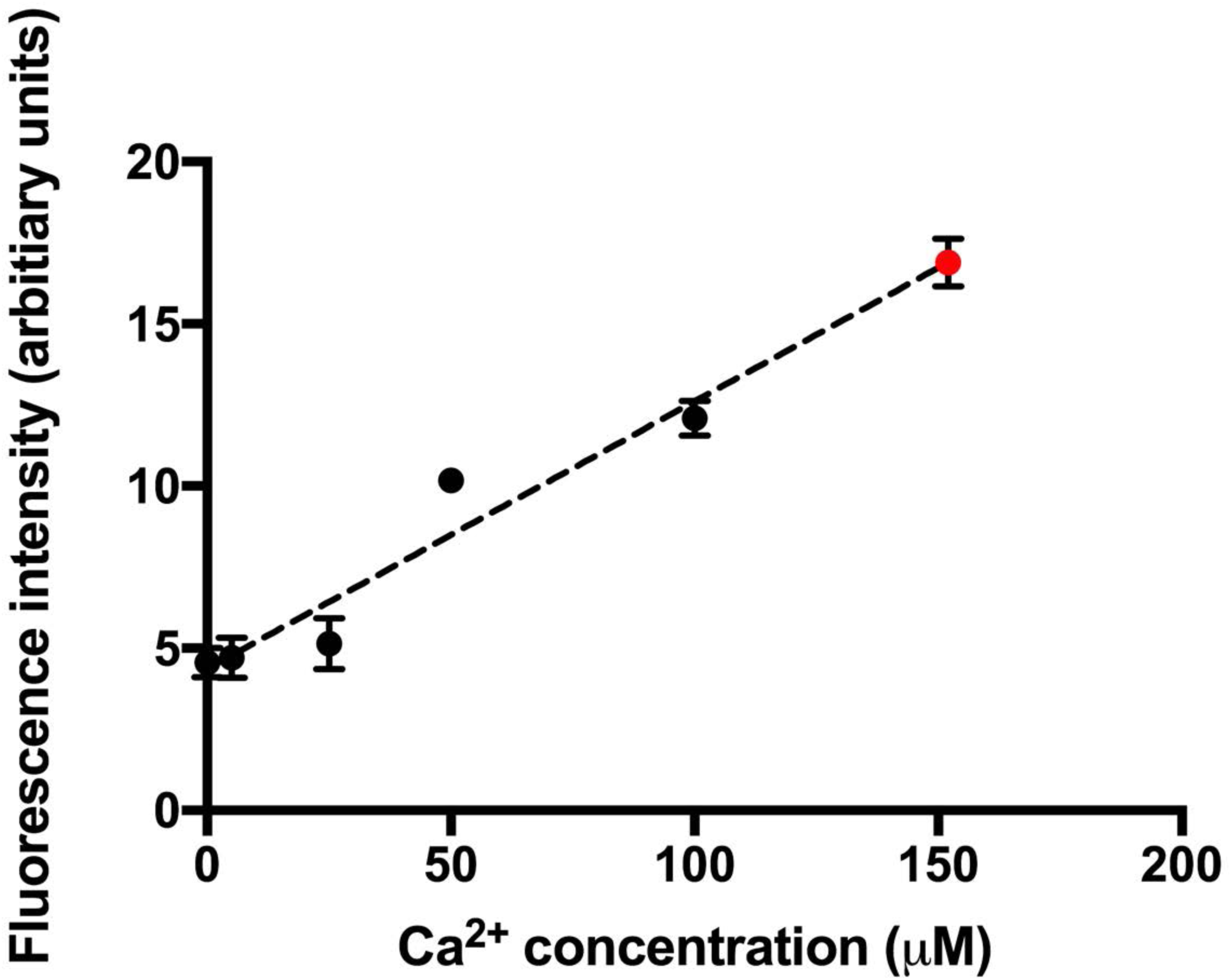
Ca^2+^ concentrations at the periphery of hair cells exceed a 100 µM. Shown is the average fluorescence intensity of hair cells with different bath concentrations of Ca^2+^. Basilar papillae from chicks were incubated in the presence of the Ca^2+^ ionophore A23187 and different concentrations of Ca^2+^ (nominally 0 µM Ca^2+^, 5 µM Ca^2+^, 25 µM Ca^2+^, 50 µM Ca^2+^ and 100 µM Ca^2+^). Fluorescence (in arbitiary units) was measured away from clusters to prevent CICR skewing our data. Measurements from 8-12 cells were averaged. Also shown (in red) is the average fluorescence intensity of Ca^2+^ clusters along the periphery of the cell in hair cells incubated in the presence of perilymph containing 1.3 mM Ca^2+^. The concentration of Ca^2+^ is in excess of 100 µM based on its fluorescence measures. Using a linear trendline, we estimate a value of 150 µM Ca^2+^. In all these measures we averaged the fluorescence from clusters at the periphery of 8-10 hair cells.

### The Slo channel clusters with both IP3 and ryanodine receptors along the periphery of the cell where they lie within a hundred nanometers of each other

Since hair cells contain high concentrations of Ca^2+^ buffer that are thought to provide significant spatiotemporal buffering, we sought to determine the localization of IP3 receptors and ryanodine receptors in hair cells in relationship to BK channels. For these experiments, we localized Slo, the BK alpha subunit, a key constituent of the BK channel complex, and the IP3 receptor using immunolabeling. We used STORM/PALM super-resolution microscopy for these experiments. As shown in **Figure 10**, both these proteins were localized along the periphery of the cell. The proteins were clustered in close proximity with one another along the periphery of the cell. In most clusters, we noted the proteins to lie in apposition in the 2D plane, or when separated to lie less than a hundred nanometers of one another. We see a similar distribution with clustering and close proximity between the Slo channel and ryanodine receptors. Here too, the proteins are closely approximated and lie within nanometers from one another along the periphery of the cell (**Figure 10**). Using the nearest neighbor algorithm, we determine that the Slo channels lie as close as 5 nm from IP3 receptors with peak distances between these proteins between 55-85 nm (**Figure 11**). Similarly, Slo channels and ryanodine receptors lie as close as 15nm apart with the peak distribution of distances between Slo and ryanodine receptors between 85 and 135 nm (**Figure 11**). These data support the presence of nanodomains within these cells, similar to that in cartwheel inhibitory interneurons of the dorsal cochlear nucleus (Irie and Trussell, 2017). The super-resolution data are concordant with our electrophysiological and Ca^2+^ imaging data.

**Figure 10.**
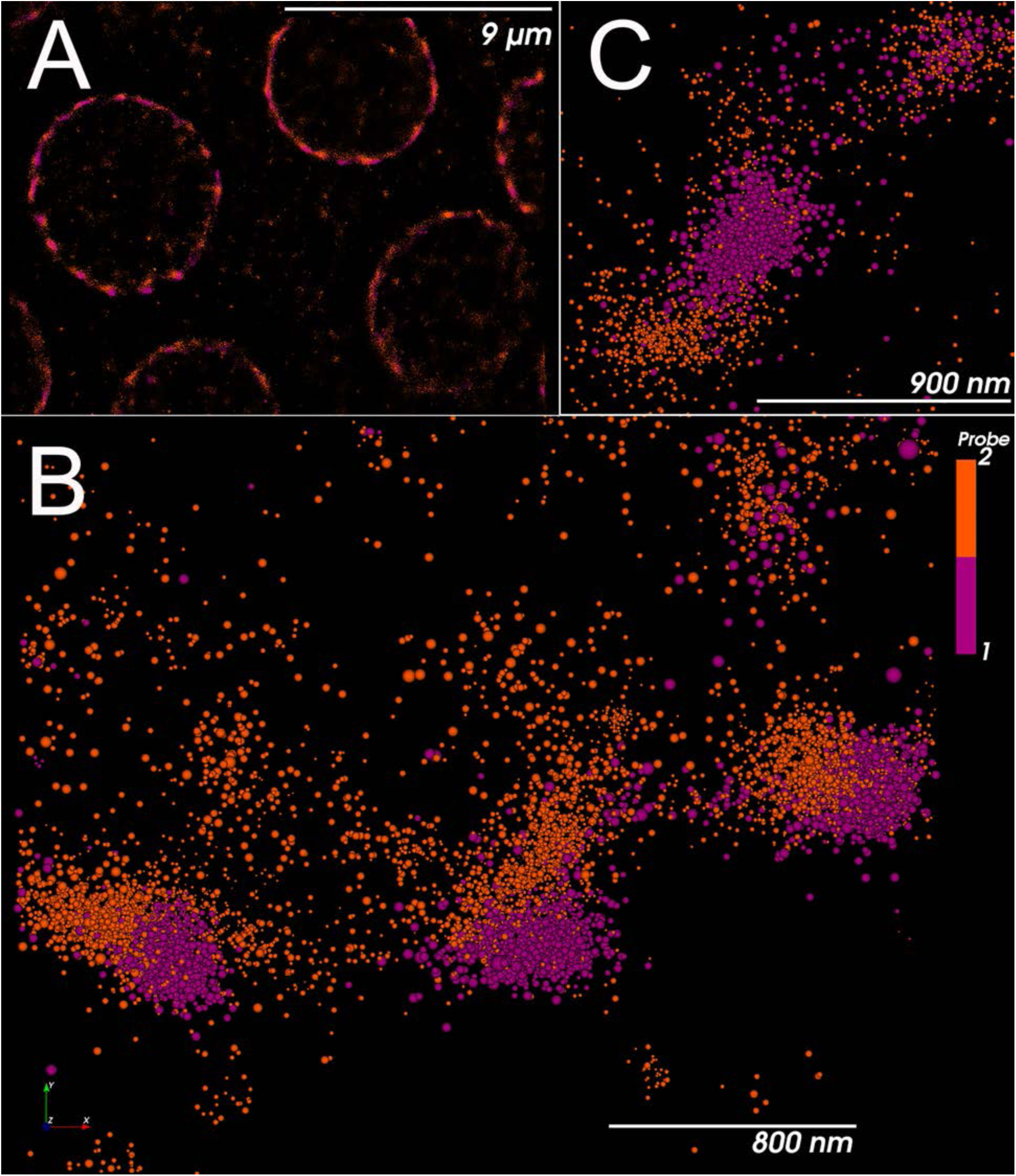
Slo channels lie in close proximity to IP3 receptors and ryanodine receptors. STORM images of individual hair cells in the basilar papilla are visualized end on from above. **A,B.** Slo channel puncta (purple) are localized in close proximity to IP3 receptors (orange) in low (A) and higher magnifications (B). Slo channel puncta lie close to the membrane with IP3 receptors arranged tangentially away from the membrane. In many instances, we could not separate the two clusters in two dimensions. Where such separation was possible, the distance between the clusters was usually less than 100 nm (see below). **C.** Slo channels (orange) are also seen in proximity to ryanodine receptors (purple). Here too, the two protein clusters were often not separable in two dimensions. Where the clusters were separable, the distance between them was usually less than 150 nm (see below).

**Figure 11.**
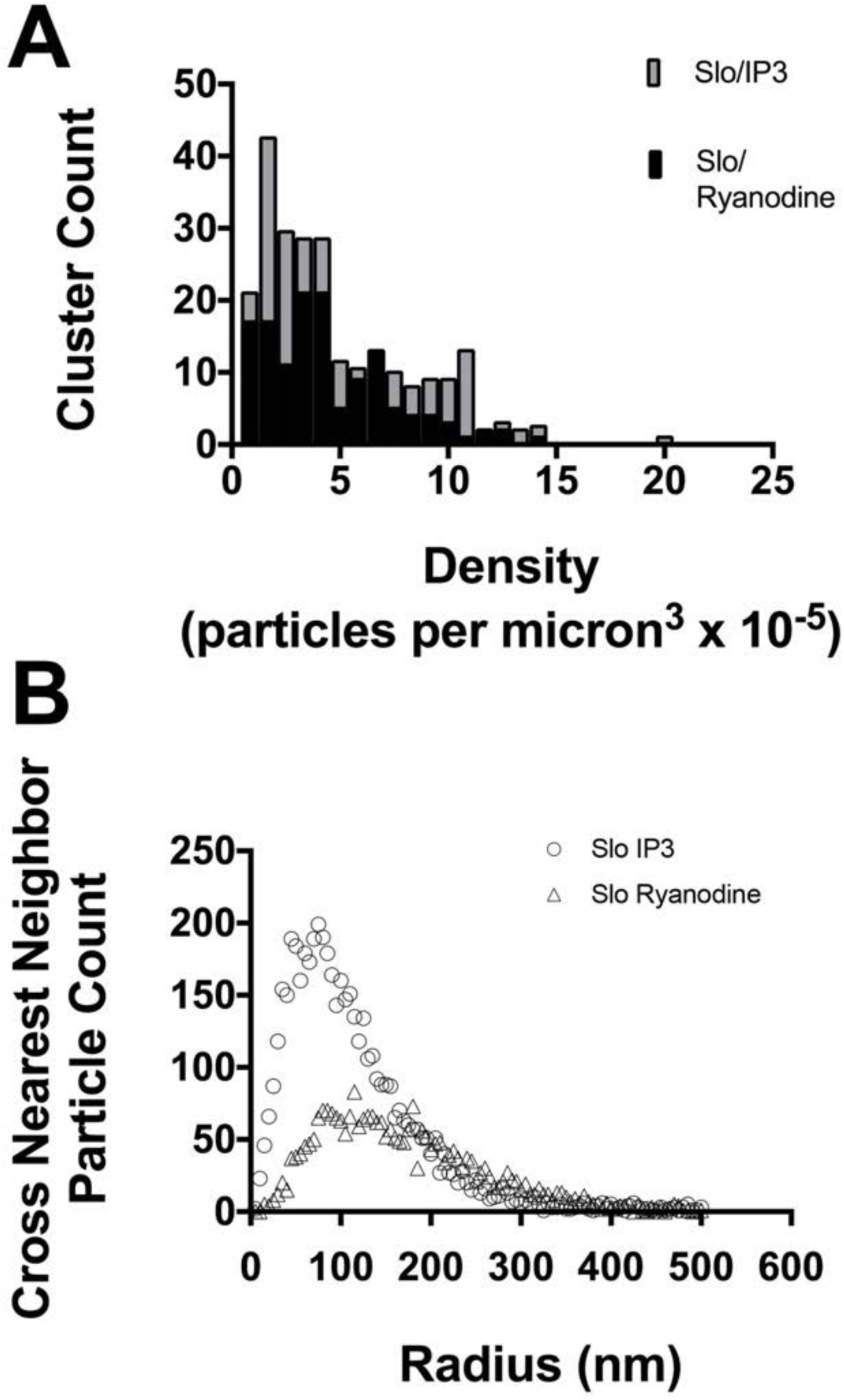
Slo, IP3 receptors, and ryanodine receptors are clustered and lie within nano-micro domains. **A**. Slo, IP3 and ryanodine receptors were clustered. Shown is the distribution in the density of particles and its variability with cluster number. The histogram of binned data shows a skewed distribution in particles. Both Slo/IP3 and Slo/ryanodine clusters show similar patterns of clustering. There was over a tenfold range in particle density in clusters. **B**. Using a Nearest Neighbor algorithm, we find a skewed distribution in the distances between Slo and IP3 receptors, and, separately, Slo and ryanodine receptors. Shown are histograms of crossed (that is centroids of Slo and IP3 particles, and, separately, centroids of Slo and ryanodine particles) nearest neighbor distances. The distances between Slo and IP3 receptors started at 5 nm, and the distances between Slo and ryanodine receptors started at 15nm. The peak in the distribution of distances between Slo and IP3 lay between 55 and 85 nm. The peak in distribution of distances between Slo and ryanodine receptors lay between 100 and 135 nm. Using a 200 nm cutoff radius, the median nearest neighbor distance was 85nm for Slo/IP3 particles, and 100nm for Slo/ryanodine receptor particles.

## Discussion

In this paper, we show for the first time the effects of CICR on BK channel function in hair cells of the chicken. We discovered the effects of CICR while exploring the effects of protein kinase A on BK channel kinetics in chick hair cells. We present three streams of data that substantiate our findings. Electrophysiological data showing a reduced current that was accompanied by a significant depolarizing shift in the I-V relationship with BAPTA compared to EGTA in nominally 0 µM Ca^2+^ suggested that higher spatial buffering afforded by BAPTA significantly limited Ca^2+^ available to activate BK channels. EGTA has a slower rate of Ca^2+^ binding (compared to BAPTA) and allows for more spatially diffuse Ca^2+^ signaling (Augustine et al., 1991; Rios and Stern, 1997; Neher, 1998). Preventing Ca^2+^ entry by blocking VGCCs and reducing extracellular Ca^2+^ both resulted in a depolarizing shift in the I-V relationship. These data suggested that entry of extracellular Ca^2+^ is important for activation of BK channels likely by raising Ca^2+^ concentrations in the vicinity of BK channels clustered and co-localized with VGCCs at the plasma membrane of hair cells. Furthermore, when using EGTA as the intracellular buffer we note an increase in the size of current with no significant change in the I-V relationship when intracellular Ca^2+^ was raised from 0 to 10 and then to 100 µM Ca^2+^. Since the use of BAPTA shifted the I-V relationship in a depolarizing direction compared to EGTA with comparable Ca^2+^ concentrations (0 µM), these data suggest that 1. local concentrations of Ca^2+^ in the presence of EGTA were saturating and 2. recruiting more BK channels are effected by higher Ca^2+^ concentrations. These data imply that BK channels are activated by two mechanisms – by entry of Ca^2+^ through VGCCs, and, farther from VGCCs, through CICR. In fact, previous confocal data have shown that majority VGCC to be in proximity to BK channels, but the majority of BK channels were spatially separated from VGCCs (Samaranayake et al., 2004). Currents in the presence of BAPTA were 20-25% of that in the presence of EGTA (both with nominally 0 µM Ca^2+^). These data argue that BK channels in approximation with VGCCs are only a fraction of BK channels in hair cells confirming prior confocal data.

When cells were perfused with forskolin we see a similar effect to that of raising pipette Ca^2+^ concentrations, namely, an increase in the size of the current. Since PKA is a well known activator of CICR and since the effects of PKA activation were dependent on pipette Ca^2^ concentration, we reason that CICR is the principle mechanism by which PKA (and raising pipette Ca^2+^ concentrations) affects BK channels. A decrease in the size of the current with specific inhibitors to both IP3 and ryanodine receptors and its prevention of forskolin induced increases in BK currents confirmed the occurrence of CICR with PKA activation.

We note differential effects of blockers of ryanodine and IP3 receptors in the presence of PKA activation. In contrast to dantrolene and forskolin, we note a greater reduction in the size of the current and a depolarizing shift in the I-V relationship when forskolin and 2APB were used in combination. One possible explanation for this result is a greater functional coupling between BK channels and IP3 receptors.

Imaging with the Ca^2+^ indicator dye Fluor-3-AM demonstrated high concentrations of Ca^2+^ along the periphery of hair cells that were clustered. The signal within these peripheral clusters was attenuated by reducing external Ca^2+^ concentration, and, separately, in the presence of inhibitors of CICR with high external Ca^2+^. Moreover, measures of Ca^2+^ concentration at the periphery of the cell using fluorescence were concordant with our electrophysiological data and supersaturating. We confirm a rise in intracellular Ca^2+^ concentrations in response to PKA activation and an attenuation or absent rise in this signal in response to PKA activation by inhibition of CICR. Together, the Ca^2+^ imaging data confirm and reinforce our electrophsiological data. Finally, using super-resolution microscopy, we confirm the presence of both IP3 and ryanodine receptors to lie within 100 nm of BK channels along the periphery of the cell.

### Nanodomains vs microdomains

In sum, our data argue for a close approximation of BK channels and CICR. Over what distances does Ca^2+^ act? This is a key question as it will determine how electrical resonance is affected by CICR in hair cells. Nanodomains have been referred to over distances of 20 nm, while microdomains refer to distances of 200 nm (Neher, 1998). Using this definition, prior work in the frog saccule suggested microdomains of Ca^2+^with synaptic release sites within 300 nm from Ca^2+^ channels (Roberts et al., 1990). While the high concentrations of local Ca^2+^ we observe and the differential responses to BAPTA and EGTA are consistent with nanodomains of Ca^2+^(Heidelberger et al., 1994; Augustine et al., 2003), our measured distances between BK channels and IP3 and ryanodine receptors extend from distances consistent with nanodomains to intermediate distances between nanodomains and microdomains. The distances between Slo and IP3 receptors, and Slo and Ryanodine receptors start at 5 nm and 15 nm, respectively. The peak in distribution of distances between BK channels and IP3 receptors was 55-85 nm. In contrast, the peak in the distribution of distances between BK channels and ryanodine receptors lay between 100-135 nm.

### How does PKA affect CICR?

How might PKA activation influence CICR in hair cells? CICR has been a most well-studied phenomenon in muscle cells, endocrine cells, and neuronal cells (Roderick et al., 2003). ITPRs acting as coincident detectors (requiring both IP3 and Ca^2+^ for activation) and ryanodine receptors that respond to Ca^2+^ have been well-studied in this context (Roderick et al., 2003). PKA increases CICR by affecting a multitude of processes in this cascade. Thus, PKA phosphorylates both ITP3Rs (ITP3R1 and ITP3R3) and ryanodine receptors 1 and 2 to increase their sensitivity to intracellular Ca^2+^, thus, increasing CICR (Islam et al., 1998,; Holz et al., 1999; Reiken et al., 2003; Dyachok and Gylfe, 2004; Wehrens et al., 2006; Taylor, 2017). Interrogation of prior published chicken Affymetrix datasets reveals that ITPR1, 2, and 3 are all detected in the basilar papilla (Frucht et al., 2011). Similarly, RYR1, 2 and 3 are all detected in the basilar papilla with no significant differential distribution of these receptors along the tonotopic axis (Itpr1, Itpr2, and Ryr1 are also expressed in mouse inner and outer hair cells) (Frucht et al., 2011; Li et al., 2018). In addition to the effects on ITPRs and ryanodine receptors, PKA also modulates phospholipase C (PLC) and couples it to receptors strengthening CICR (Liu and Simon, 1996). Finally, PKA has been shown to phosphorylate Cav1.3 increasing its conductance (Mahapatra et al., 2012). Our data showing a block of PKA effects by inhibiting CICR suggests that the effects of PKA are predominantly effected through CICR and not through effects on Cav 1.3.

### Effects on electrical resonance

What effects might CICR have on electrical resonance? While we did not explore the effects of PKA activation and CICR on electrical resonance, our initial impetus to studying PKA effects on BK currents was to explore how varying BK channel kinetics occur in hair cells along the tonotopic axis. Varying BK channel kinetics is the principal mechanism for frequency tuning and electrical resonance in the turtle (Art et al., 1995). There is data showing a similar mechanism operates in the chick (Fuchs et al., 1988; Fettiplace and Fuchs, 1999).

The close proximity of IP3 and ryanodine receptors to BK channels could extend the distance of Ca^2+^ signaling, thereby extending its operational range. This could explain the larger currents observed with CICR from an increasing number of BK channels activated farther from the site of Ca^2+^ entry. On the other hand, CICR could simultaneously potentially attenuate the temporal fidelity of electrical resonance from the increased duration of feedback inhibition. Other variables that affect the sharpness of the negative feedback loop between voltage-gated Ca^2+^ channels and BK channels are the distance between these two channels (estimated to be 50nm), the number of Ca^2+^ and BK channels at these local clusters, and the effective buffering of the many different native Ca^2+^ buffers (Roberts, 1993, 1994; Wu et al., 1995). Previous modeling experiments in the turtle have incorporated these variables and produced a reasonable approximation to experimental data (Wu et al., 1996). Our data showing CICR and its effects on BK channels would necessitate a rethinking of these models, particularly if CICR is shown to operate in the turtle, as well. With data showing complex control and kinetics of CICR in other systems, our data in turn point to increasingly complex control of electrical resonance (Dyachok and Gylfe, 2004).

Our data with whole-cell recordings showed no change in the voltage dependence of BK currents with 100 µM, 10 µM and nominally 0 µM intracellular Ca^2+^ when EGTA was used as a buffer. On face value, these data could be taken to imply that the concentration of Ca^2+^ in proximity to BK channels are a 100 µM even when the pipette Ca^2+^ concentration was nominally 0 µM Ca^2+^ owing to the limited spatial buffering of EGTA with (extracellular) Ca^2+^ entry occurring through voltage-gated channels. Our measurements of Ca^2+^ concentration as exceeding 100 µM, albeit in the native state where buffering capacity is not precisely defined (hair cells contain mM concentrations of Ca^2+^ buffer), are consistent with this possibility.

The high local concentration of Ca^2+^ is also consistent with electrophysiological measurements of Ca^2+^ sensitivity of BK channels in these hair cells. Duncan et al., using excised patch recordings, demonstrated a half-maximal Ca^2+^ sensitivity of 0.1 – 5 µM Ca^2+^ at +50mV (the voltage at which our measurements of currents in whole-cell recordings were robust) (Duncan and Fuchs, 2003). Close to the resting membrane potential of hair cells (−50mV), these authors determine a half-maximal Ca^2+^ concentration of 5-100 µM (although at the frequency location we used – 20-30% from the apical end-the concentration was closer to 5-50 µM).

The high local concentration of Ca^2+^ in proximity to BK channels from CICR could also explain the seeming discrepancy in in-vivo findings of minimal differences in Ca^2+^ sensitivity compared to the large differences in Ca^2+^ sensitivity observed when splice variants were expressed in heterologous systems (Art et al., 1995; Jones et al., 1999; Ramanathan et al., 1999; Ramanathan et al., 2000; Duncan and Fuchs, 2003). Excised patches used to determine sensitivity to Ca^2+^ would presumably also contain IP3 and ryanodine receptors that in turn would induce the local release of Ca^2+^.

### Other implications of CICR in hair cells

Synaptic vesicle release is the other mechanism that is affected by Ca^2+^ entry into the base of hair cells. How might CICR affect synaptic release in hair cells? Increasing BK current size with no effects of its voltage sensitivity points to a spatially enlarging Ca^2+^ signal with CICR in chick hair cells. A spatially extended Ca^2+^ signal could also increase the release of synaptic vesicles. In fact, such a possibility has been suggested by experimental data in turtles (with corroborating data from mice and rats). Real-time measurements of synaptic release (capacitance measures) and Ca^2+^ imaging show a linear and supralinear relationship between cell Ca^2+^ and synaptic release in inner hair cells (Schnee et al., 2011). The supralinear release was correlated with an additional intracellular source of Ca^2+^ and was speculated to be responsible for recruiting the reserve pool of vesicles.

In conclusion, we show that CICR resides in hair cells in close proximity to BK channels. We provide electrophysiological, confocal Ca^2+^ imaging, and super-resolution fluorescence immunolocalization data to support our conclusion that CICR likely plays an important role in hair cell function. These data require a rethinking of the physiological and molecular mechanisms of electrical resonance and synaptic vesicle release.

